# High-efficiency genomic mapping of chromatin-associated targets with CUT&RUN

**DOI:** 10.1101/2024.12.03.626419

**Authors:** Tessa M. Firestone, Bryan J. Venters, Katherine Novitzky, Liz Marie Albertorio-Sáez, Courtney A. Barnes, Karlie N. Fedder-Semmes, Nathan W. Hall, Allison R. Hickman, Mark Kaderli, Carolina Lin Windham, Matthew R. Marunde, Danielle N. Maryanski, Kelsey Noll, Leslie Shannon, Jennifer Spengler, Martis W. Cowles, Zu-Wen Sun, Michael-Christopher Keogh, Andrea L. Johnstone, Ellen N. Weinzapfel, Lu Sun

## Abstract

The precise regulation of chromatin composition is critical to gene expression and cellular identity, and thus a key component in development and disease. Robust assays to study chromatin features, including histone post-translational modifications (PTMs) and chromatin-associated proteins (*e.g.*, transcription factors or PTM readers), are essential to understand their function and identify novel therapeutic strategies. To this end, Cleavage Under Targets and Release Using Nuclease (CUT&RUN) has emerged as a powerful tool for high-resolution epigenomic profiling. The approach has been successfully applied to numerous cell and tissue types, informing on target genomic distribution with unprecedented sensitivity and throughput. Here, we provide a detailed CUT&RUN protocol from sample collection through data analysis, including best practices and defined controls to ensure specific, efficient, and robust target profiling.

## INTRODUCTION

### 1. Chromatin mapping approaches

Chromatin refers to the structural organization of the genome in eukaryotic cells, and is central to gene expression, cell division, and DNA repair [1, 2]. This protein-nucleic acid assemblage is regulated by diverse modifications (including histone post-translational modifications (PTMs) [3, 4] DNA methylation [5]), chromatin-associated proteins (CAPS; including transcription factors [6], chromatin readers / editors [7], and remodeling enzymes [8]). The dysregulation of chromatin features can operate in concert with genetic mutations and environmental factors [9–12] to alter gene expression profiles and drive cancer development [13–19]. As such, mapping the genomic localization of histone PTMs and CAPs can reveal relationships between the chromatin landscape and disease phenotypes. To this end, researchers have historically relied on <u>Ch</u>romatin Immuno<u>P</u>recipitation (ChIP)-based approaches, most notably *ChIP-seq*. This has provided fundamental insights to chromatin structure / function and advanced the application of epigenomics across many biomedical fields. However, the inherent limitations of *ChIP-seq*, which requires high cell numbers and deep sequencing to overcome high background and technical fragility, have precluded its widespread application to drug research and clinical studies. Further, *ChIP-seq* can suffer from technical artifacts arising from experimental manipulations, including heavy crosslinking and its reversal. Assays with improved sensitivity, robustness, and scalability are needed to better inform (pre)clinical research and identify novel diagnostic and therapeutic opportunities.

To address these needs, a new epigenomic mapping method has recently emerged: **<u>C</u>**leavage **<u>U</u>**nder **<u>T</u>**argets **<u>& R</u>**elease **<u>U</u>**sing **<u>N</u>**uclease (CUT&RUN) [20, 21]. In the approach [**Figure 1**], cells or nuclei are immobilized to Concanavalin A (ConA) coated magnetic beads, permeabilized, and incubated with target-specific antibody. A pAG-MNase (protein A-protein G-Micrococcal Nuclease) fusion is added to selectively cleave and release antibody-bound chromatin to the supernatant. Bead-coupled cells remain on the magnet, while CUT&RUN enriched DNA is purified from the supernatant and prepared for multiplexed next-generation sequencing (NGS). Finally, CUT&RUN reaction libraries are sequenced to a depth of 5-7 million total reads (assuming a human genome), which provides sufficient coverage for high-resolution genomic mapping.

**Figure 1.**
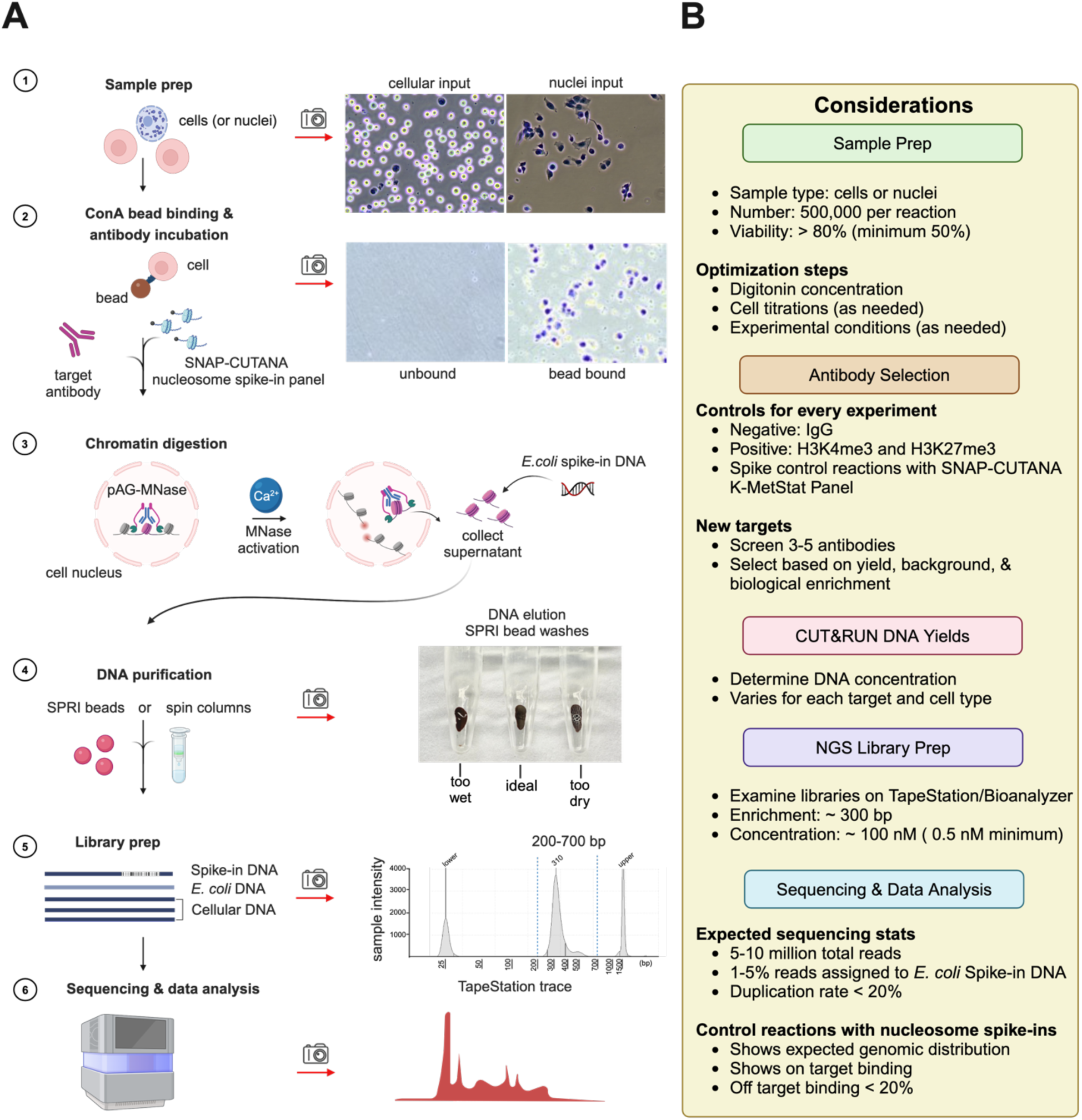
The CUT&RUN workflow^1^. **(A)** Schematic of a representative CUT&RUN experiment. Camera icons depict expected observations at key quality control (QC) steps. **(1)** The quality of monodispersed cells or nuclei is determined by Trypan Blue staining, where the dye is excluded by native ‘viable’ cells (K562 shown), but enters permeabilized cells or isolated nuclei (MCF7 shown) **[NOTES 17-19]**. **(2)** Cells or nuclei are immobilized to ConA magnetic beads. Efficient bead binding is confirmed by examination of bound and unbound fractions under a microscope (HEK293 shown). The bead slurry is then aliquoted to 8-strip tubes for CUT&RUN. If applicable for the reaction target, SNAP-CUTANA^TM^ nucleosome spike-ins (EpiCypher) are added **[NOTES 1, 8-10]**. Target-specific antibody **[NOTES 2-10]** is added and reactions incubated overnight at 4°C. **(3)** pAG-MNase is added, and activated by further addition of Ca^2+^, leading to selective cleavage and diffusion of antibody-labelled chromatin fragments to the supernatant. Reactions are supplemented with exogenous E. coli spike-in DNA to monitor to control for input and support normalization after NGS **[NOTES 40]**. **(4)** Bead-bound cells are removed on a magnet and target-enriched DNA purified from the supernatant. If using SPRI beads, avoid over-drying (they should retain a damp dark brown color) before elution. **(5)** CUT&RUN DNA is quantified by fluorometric assays (e.g., Qubit^TM^, Invitrogen^TM^) and used to prepare sequencing libraries. Purified libraries are examined on a TapeStation or Bioanalyzer (Agilent Technologies) to determine molarity and confirm enrichment of mononucleosome-sized fragments (∼300 bp) **[NOTES 48-52]**. **(6)** Pooled CUT&RUN libraries are sequenced (e.g., Illumina), aiming for 5-10 million total reads per reaction (assuming a human genome), and data analyzed **[NOTES 53-59]**. **(B)** Important considerations at each step to ensure a successful CUT&RUN experiment.

Importantly, CUT&RUN offers dramatic improvements in throughput, sensitivity and cost over *ChIP-seq* [22–24]. The high Signal-to-Noise (S:N) of the new approach is achieved by fewer, lower stringency washes and the direct partition of target-enriched chromatin fragments (signal) from bead-coupled cells (noise). This enables reduced cell numbers (5-500 k per reaction) and sequencing depth (5-7 M total reads per reaction assuming a human sample), compared to the >1 M cells and >30 M reads needed for *ChIP-seq* [25, 26]. To further improve throughput and reproducibility at reduced cost, this CUT&RUN protocol is adapted for 8-strip tubes and multichannel pipettors. In our experience, the practiced user can process 24-48 reactions over a two- to three-day procedure. This throughput supports the inclusion of control reactions for each biological sample and encourages deeper experimental design: allowing the user to examine more replicates, time points, inhibitor doses, or additional targets.

Studies using CUT&RUN have delivered high-quality datasets for multiple histone PTMs and CAPs [27–31] from a diversity of sample inputs, including cell lines, primary tissues, and model organisms [32–48]. To illustrate its feasibility, we used CUT&RUN to study cellular responses after inhibition of the chromatin remodeler SMARCA2 (BRM) [**Figure 2**]. BRM014 allosterically inhibits SMARCA2 catalytic activity [49] without impacting global abundance [50, 51]; while the AU-15530 and ACBI1 <u>pro</u>teolysis <u>ta</u>rgeting <u>c</u>himeras (PROTACs) selectively engage and degrade the ATPase [51, 52], leading to signal loss on chromatin [**Figure 2B**]. This highlights a major advantage of CUT&RUN for parallelized drug screening studies: the workflow can be easily scaled for additional compounds, doses or cell types.

**Figure 2.**
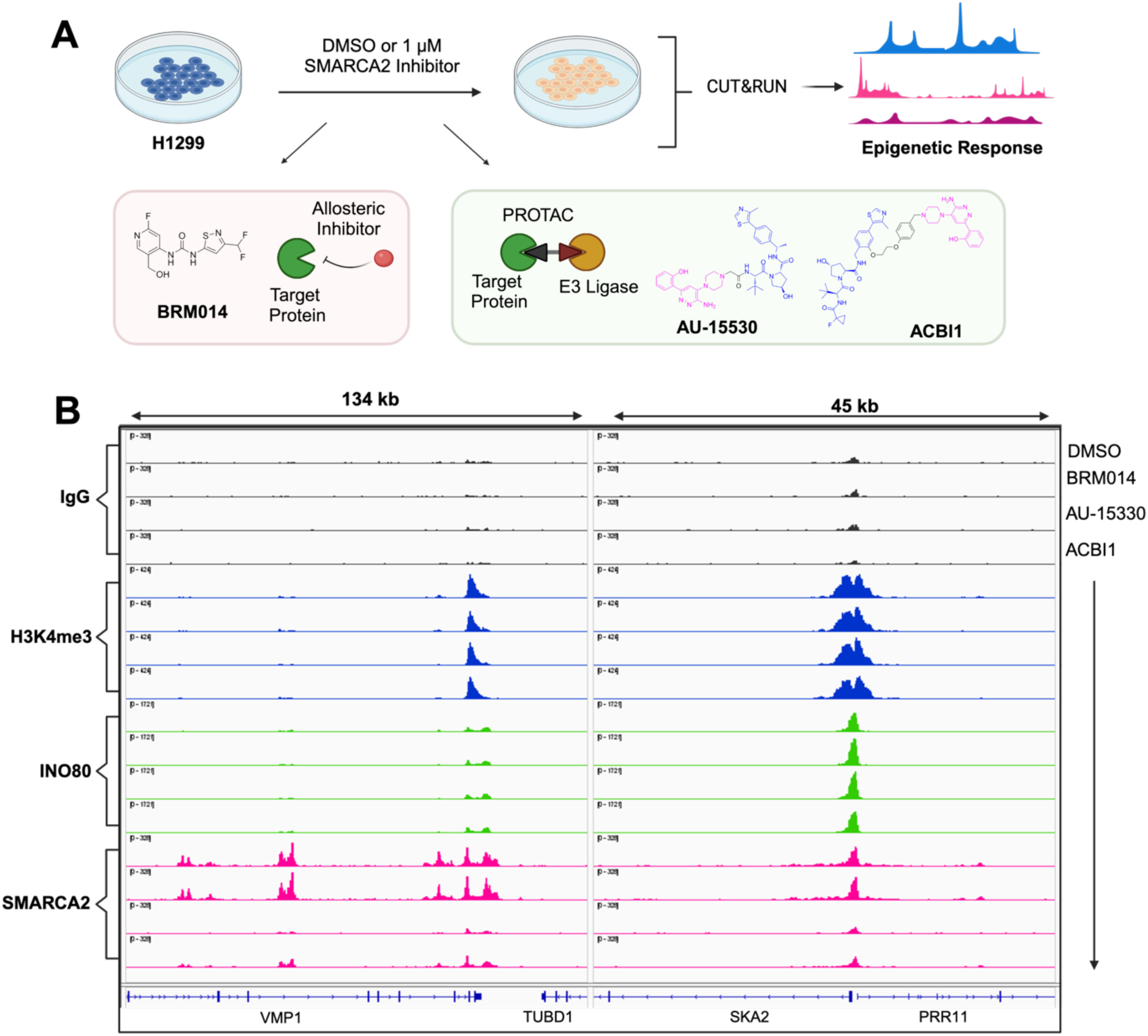
Application of CUT&RUN to study different classes of SMARCA2/BRM inhibitors^2^. **(A)** Design of a parallelized CUT&RUN experiment testing various inhibitors of the SMARCA2 (BRM) chromatin remodeling ATPase. H1299 cells were treated for four hours with BRM014 [49], AU-15330 [51] or ACBI1[52] (all 1 µM); or DMSO vehicle control (0.01 %). Cells were lightly cross-linked (0.1% formaldehyde for 1 min) **[NOTE 16]** and processed by CUT&RUN. **(B)** Browser tracks (IGV) at representative genomic locations [78] after indicated treatment, showing signal for IgG (negative control), H3K4me3 (control; active promoters), INO80 (control; a non-targeted remodeler), and SMARCA2 (inhibitor target). BRM014 or DMSO had no impact on SMARCA2 or INO80 enrichment, while the PROTACs (AU-15530 and ACBI1) reduced SMARCA2 but not INO80 binding to chromatin, confirming their selective degrader mechanism of action. RPKM normalized reads are group-scaled by target and aligned to human genome build hg38.

This chapter provides a detailed CUT&RUN protocol optimized for high throughput and S:N at reduced sequence depth [**Figure 1**]. We discuss the criteria for identifying the best antibodies for a given target (*e.g.*, histone PTM or CAP); methods to prepare CUT&RUN-ready samples; how to navigate optimization steps; and considerations for data analysis. We recommend all experiments include reactions with control antibodies and/or spike-ins that provide insight to various aspects of the CUT&RUN workflow, including sample quality, cell permeabilization, antibody performance, pAG-MNase activity and library preparation. This includes reactions controlled by DNA-barcoded nucleosomes (*e.g.*, *EpiCypher* SNAP-CUTANA^™^ K-MetStat or epitope tag panels) **[NOTE 1]** that can be seamlessly integrated to the workflow and greatly increase confidence in the resulting CUT&RUN datasets. Finally, we include sections on library preparation for *Illumina* NGS with considerations for sequencing depth, quality metrics, and data analysis.

## 2. MATERIALS

### 2.1 Equipment

Aluminum Block compatible with strip tubes: *Thermo Fisher Scientific* NC9820161 (optional)

Automated capillary electrophoresis instrument: *e.g.*, *Agilent* Bioanalyzer or *Agilent* TapeStation

Cell Counter: *Invitrogen* AMQAX2000

8-channel multichannel pipettes: *VWR* 76169-250

Fluorometer: e.g., *Invitrogen* Qubit 4 Q33238

Hemocytometer slides: *Sigma-Aldrich* Z359629

Illumina sequencer: *e.g.*, NextSeq1000/2000

Incubator: *Millipore Sigma* Z763314

Magnetic rack for 8-strips: *EpiCypher* 10-0008

Magnetic rack for 1.5 mL tubes: *EpiCypher* 10-0012

Microcentrifuge: *Thermo Fisher Scientific* 75002435

Minicentrifuge: Thermo Fisher Scientific 75004086

Microscope: *Nikon* Eclipse Ts2

Nutator: *VWR* 82007-202

Thermocycler with heated lid: *e.g.*, *BioRad*, *Applied Biosystems*, *Eppendorf* (or comparable)

Vortex Mixer: Scientific Industries SI-0236

### 2.2 Consumables and reagents (* optional)

Automated capillary electrophoresis consumables (e.g., *Agilent* High Sensitivity DNA Analysis Kit 5067-4626; or D1000 ScreenTape 5067-5582 and D1000 Reagents 5067-5583)

CaCl_2_ (e.g., Sigma-Aldrich C1016)

Cell counter slides (e.g., *Invitrogen* C10228)

Cell culture and appropriate growth medium

ConA Beads (*e.g.*, *EpiCypher* 21-1401)

Control Antibodies

- IgG Negative Control (*e.g.*, *EpiCypher* 13-0042)
- H3K4me3 Positive Control (*e.g.*, *EpiCypher* 13-0060)
- H3K27me3 Positive Control (*e.g.*, *EpiCypher* 13-0055)
- H3K27ac Positive Control*, labile PTM (*e.g.*, *EpiCypher* 13-0059)

Digitonin (5% stock in DMSO) (*e.g.*, *Millipore Sigma* 300410)

DMSO (e.g., *Sigma-Aldrich* D8418)

DNA purification kit (*e.g.*, *EpiCypher* 14-0052)

*E. coli* Spike-in DNA (*e.g.*, EpiCypher 18-1401)

EDTA (0.5 M pH 8) (*e.g.*, *Sigma-Aldrich* E5134)

EGTA (0.5 M pH 8) (*e.g.*, *Sigma-Aldrich* E3889)

Formaldehyde* (37% w/v) (*e.g.*, *Sigma-Aldrich* 252549)

Fluorometer reagents (*e.g.*, *Invitrogen* 1X dsDNA HS Kit Q33230)

Glycerol (*e.g.*, *Sigma-Aldrich* G5516)

Glycine* (2.5M) (*e.g.*, *Sigma-Aldrich* 50046)

Glycogen (20 mg/mL) (*e.g.*, *Sigma-Aldrich, Roche* 10930193001)

HEPES (e.g., *Sigma-Aldrich* H3375)

HEPES (0.5M, pH 7.9) [autoclaved or filter sterilized] [stocks for **2.3**]
HEPES (0.5M, pH 7.5) [autoclaved or filter sterilized] [stocks for **2.3**]

KCl (e.g., *Sigma-Aldrich* P3911)

Library prep kit(s) (*e.g.*, *EpiCypher* 14-1001 or 14-1002 [each provides different 48x multiplex barcode sets] or NEB NEBNext Ultra II DNA Library Kit E7645, E7600S, & E7780S)

Low-retention filter pipette tips: any vendor

MnCl_2_ (*e.g.*, *Sigma-Aldrich* 203734)

Molecular biology grade water (*e.g.*, *Corning* 46-000-CM)

Multichannel reagent reservoirs: *Thermo Fisher Scientific* 14-387-072

NaCl (*e.g.*, *Sigma-Aldrich* S5150-1L)

Nuclei Extraction Buffer* (*e.g.*, *EpiCypher* 21-1026)

100% Ethanol (200 proof) (*e.g.*, *Millipore Sigma* EX0276)

1.5, 15, 50 mL tubes: any vendor

1X PBS* (calcium and magnesium free) (*e.g.*, *Cytiva* SH30256.01)

pAG-MNase (*e.g.*, *EpiCypher* 15-1016)

Propidium Iodide* (*e.g.*, *Thermo Fisher Scientific* BMS500PI)

Protease Inhibitor (*e.g.*, *Roche* 11873580001)

RNase A (10mg/mL) (*e.g.*, *Thermo Fisher Scientific* EN0531)

SDS (10% solution): any vendor

SNAP-CUTANA Nucleosome Panels* **[NOTE 1]**

- *e.g.*, K-MetStat (*EpiCypher* 19-1002)
- *e.g.*, HA Tag (*EpiCypher* 19-5002)
- *e.g.*, DYKDDDDK Tag (*EpiCypher* 19-5001)

Spermidine trihydrochloride (molecular weight 254.63) (*e.g.*, *Sigma-Aldrich* S2501)

- Spermidine (1M) [stocks for **2.3**]

SPRIselect Bead-Based Reagent (e.g., *Beckman Coulter* B23318)

Test antibodies **[NOTES 2-10]**

Tris Base (*e.g.*, *Genesee Scientific* 18-145)

- Tris (1M, pH 8.0) [autoclaved or filter sterilized] [stocks for **2.3**]
- Tris (1M, pH 8.5) [autoclaved or filter sterilized] [stocks for **2.3**]

Triton X-100 (10% solution) (*e.g.*, *Sigma-Aldrich* T8787)

Trypan Blue (0.4%) (*e.g.*, *Invitrogen* T10282)

Trypsin* (0.05%) (*e.g.*, *Cytiva* SH30236.01)

Tween 20 (*e.g.*, *VWR* 9005-64-5)

0.2 mL 8-strip tubes with attached flat caps: *EpiCypher* 10-0009

### 2.3 Buffer Recipes (* optional)

All buffers are made with molecular biology grade water.

**Wash**, **Antibody**, and **Nuclei Extraction Buffers** are made fresh (same day) for each experiment by adding spermidine, digitonin or protease inhibitor (as individually required) to respective precursor buffers. **Cell Permeabilization Buffer** can be made on Day 1 and stored at 4°C for use on Day 2. **Stop** and **Bead Activation Buffers** can be filter sterilized and stored at 4°C for ∼ 6 months.

**5% Digitonin:** Prepare 5% (w/v) digitonin in DMSO. Stable for 6 months at -20°C.

**2.5 M Glycine *:** Dissolve 9.4g in 50 mL molecular grade water. Filter sterilize. Stable at room temperature (RT°C) for 6 months.

**25X Protease Inhibitor:** Dissolve 1 protease inhibitor tablet in 2 mL water. Stable for 12 weeks at -20°C.

**1M Spermidine:** Dissolve 1 g Spermidine trihydrochloride (MW 254.63) in 3.93 mL water. Stable at -20°C for 6 months store in single-use aliquots. **[NOTE 11]**

**Bead Activation Buffer**: 20 mM HEPES pH 7.9, 10 mM KCl, 1 mM CaCl_2_, 1 mM MnCl_2_. Filter sterilize. Stable at 4°C for 6 months.

**Nuclei Extraction Buffer**: 20 mM HEPES pH 7.9 (pH with KOH), 10 mM KCl, 0.10% Triton X-100, 20% glycerol. Filter sterilize. Stable at 4°C for 6 months. On **Day 1** of each experiment supplement with 1X protease inhibitor and 0.5 mM spermidine and store at 4°C. Discard after **1 day.**

**Pre-wash Buffer**: 20 mM HEPES pH 7.5, 150 mM NaCl. Filter sterilize. Stable at 4°C for 6 months.

**Wash Buffer**: **Pre-wash Buffer** freshly supplemented with 1X protease inhibitor and 0.5 mM spermidine. Use within **Day 1** of each experiment. **[NOTE 12]**.

**Cell Permeabilization Buffer**: **Wash Buffer** freshly supplemented with digitonin (to 0.01%) on **Day 1** of each experiment. Stable at 4°C overnight. See (**4.2**) for digitonin optimization.

**Antibody Buffer**: **Cell Permeabilization Buffer** freshly supplemented with EDTA pH 8.0 (to 2 mM). Use within **Day 1** of each experiment. (**4.3.3**).

**Stop Buffer**: 340 mM NaCl, 20 mM EDTA pH 8.0, 4 mM EGTA pH 8.0, 50 µg/mL RNase A, 50 µg/mL glycogen. Filter sterilize. Stable at 4°C for 6 months.

**0.1X TE Buffer**: 1 mM Tris pH 8, 0.1 mM EDTA pH 8.0. Filter sterilize. Stable at RT°C for 6 months.

**Sequencing Buffer**: Illumina RSB with Tween 20, Cat # 20050639. Stable at 4°C for 6 months.

## 3. Antibody selection

Delivering the full high-sensitivity / low background potential of CUT&RUN requires antibodies of high specificity and efficiency. The presumptuous adoption of unsuitable antibodies is a major and widespread issue in biomedical research [53–55]. This problem is especially relevant in genomic mapping assays, where antibodies validated by ELISA or immunoblot are often assumed to perform similarly in the very different biophysical context of chromatin [56]. Such surrogate validations have caused great harm, with many of the resulting ‘approved’ histone PTM antibodies showing high cross-reactivity and/or poor efficiency in ChIP with spike-in nucleosome standards [56–58]. Further, even appropriately ChIP-validated antibodies are not guaranteed to similarly perform in CUT&RUN **[NOTE 2]**, demonstrating the necessity for assay-specific validation. Finally, antibodies as biological products often display lot-to-lot variability: polyclonals (pAbs) >> monoclonals (mAbs), but still experimentally observed for the latter [55, 59].

Encompassing all the above: for accurate and robust CUT&RUN, antibodies need to be thoroughly vetted, directly validated for in-assay performance, and continuously monitored. Reagent efficiency and specificity metrics are particularly critical. **High efficiency** is the greatest possible capture of on-target events and can be explored by reducing cell numbers and/or using samples with reduced target abundance. **High specificity** refers to low cross-reactivity, and can be assayed by target deletion / depletion or DNA-barcoded nucleosome spike-ins. Here, we describe how to select histone PTM and CAP antibodies that generate high-quality CUT&RUN profiles [**Figure 3**].

### 3.1 Obtain antibodies

1. Source three to five antibodies to the PTM or CAP of interest. Each should be distinct (*i.e.*, not the same reagent distributed by multiple entities), which can be achieved by ensuring different immunogens, host species or mAb clone identifiers **[NOTES 3-5]**.

### 3.2 Plan your experiment

2. Start with 500 k cells per reaction **[NOTE 6]** in a native or lightly fixed form **[NOTES 13-16]**. When establishing the CUT&RUN approach or testing novel detection reagents it is often most convenient to use an easily grown cell line wherein the target is known to be expressed and bind chromatin **[NOTES 6-7]**.
3. Antibodies to histone PTMs or epitope tags can be validated and monitored *in situ* using SNAP-CUTANA DNA-barcoded nucleosome spike-ins **[NOTES 1, 8]**.
4. Include parallel reactions with negative (IgG) and positive (H3K4me3, H3K27me3) control antibodies and appropriate SNAP-CUTANA K-MetStat nucleosome spike-ins **[NOTES 9-10].**
5. Relative efficiency for each antibody can be determined by titrating cell numbers with capability for the intended study identified by bracketing 500 k (the initial recommendation) to the likely available input (if precious or rare) [*e.g.*, **Figure 3C** and **NOTE 6**].

### 3.3 Antibody performance: analysis and comparison

Confirm positive and negative controls performed as expected. Troubleshoot the workflow as needed **[NOTES 9-10]**. For novel targets, select the antibody with best balance of CUT&RUN yields, S:N and distinct genomic enrichment [**Figure 3D**].

6. Antibodies with acceptable <u>specificity</u> to PTMs or epitope-tags should show <20% cross-reactivity to SNAP-CUTANA spike-in panels [**Figures 3A-B**].
7. Antibodies with acceptable <u>efficiency</u> can be identified by maintaining peak enrichment at reduced sample input [**Figure 3C**]. Lower efficiency reagents, in contrast, will manifest decreased peak number, lower FRiP scores and degraded peak enrichment; eventually reaching IgG-like profiles.
8. Success with an ‘antibody screening to a novel target’ campaign can be determined by various considerations [*e.g.*, **Figure 3D** and **NOTE 10**]

a. Peak enrichment metrics indicate high Signal:Noise (S:N).
b. Two or more independent reagents yield highly similar peak locations.
c. The genomic distribution of peaks is consistent with reported target function or correlated epigenetic marks (not being blind to a reported function being wrong).

## 4. METHOD

### 4.1 Sample Prep

Perhaps the major variables to CUT&RUN experimental success are the antibody (**3**) and sample. The latter is impacted by a range of considerations:

1. Source and processing (*e.g.,* tissues, primary cells or lines; cells or nuclei; fresh, frozen or lightly cross-linked) **[NOTES 13-16** and **21]**
2. Cell quality (*e.g.,* material from a necrotic tissue, dying culture or after poor cryo-preservation will show increased background) **[NOTES 17-19]**
3. Cell numbers (starting recommendation is 500 k cells/reaction; though as above, high quality antibodies will often give success from fewer) **[NOTES 6** and **17-18]**
4. Effective cell permeabilization [*e.g.*, by Digitonin (**4.2**)] and/or nuclei isolation [**NOTES 13-15** and **23**]

This section describes sample processing workflows to generate reliable CUT&RUN data from native or lightly-fixed cells or nuclei [**Figure 4**]. We have found that data quality for many targets using cryopreserved material is indistinguishable from fresh samples (and the former greatly increases convenience). Nuclei extraction is recommended when performing CUT&RUN on immune cells and often required when working with tissues **[NOTES 13-15].**

Of note, the heavy cross-linking strategies of *ChIP-seq* (*e.g.*, 1% formaldehyde for 10 min) are not suitable for CUT&RUN since they dramatically reduce yields and S:N **[NOTE 16]**. However, CUT&RUN is compatible with frozen and/or lightly cross-linked material (0.1% formaldehyde for 1 min). Instances where light cross-linking is particularly helpful (or even essential) include stabilizing labile PTMs (*e.g.*, acetylation, ubiquitination or phosphorylation) and after disruptive treatment time courses.

Optimal samples are monodispersed cells (or nuclei) with normal morphology and minimal cell lysis debris. Low starting cell viability and/or poor morphology generally increases assay background. We recommend checking samples by microscopy at three points during early CUT&RUN steps: **(i)** initial cell harvest; **(ii)** after cells have been washed in CUT&RUN **Wash Buffer** (or after nuclei have been extracted), just prior to bead binding; and **(iii)** after binding to ConA beads. Performing these QC steps will identify problem samples and contribute to assay success.

The recommended starting point is 500 k cells (or nuclei) per CUT&RUN reaction. When collecting cells, process a minimum of 20% excess to account for QC checks and sample loss during wash steps. While high performance antibodies will yield robust S:N from fewer cells **[NOTE 6]**, it is recommended to first validate reagents at standard cell numbers before titrating to lower input levels.

#### 4.1.1 Counting cells and examining sample integrity

Cell quality / membrane integrity can be checked at CUT&RUN **Day 1** by Trypan Blue exclusion. Alternatively, the membrane impermeant fluorescent DNA stain propidium iodide (PI; used as per manufacturer’s recommendations) improves accuracy with compatible automated cell counters, especially when using samples from trypsinized adherent cell lines or primary tissue that are often accompanied by higher debris levels. Either stain can be used to examine samples at initial processing (**4.1.2**), to monitor effective permeabilization with Digitonin (**4.2**), after resuspending in CUT&RUN **Wash Buffer** (**4.3.2**), or to confirm nuclei extraction **[NOTES 17-19]**.

1. Add 10 µL of Trypan Blue (0.4%) to 10 µL cells and gently pipette 10 times to mix **[NOTE 18]**.
2. Immediately transfer 10 µL of mix to a hemocytometer or automated cell counter.
3. View under brightfield microscope or in dedicated cell counter. Dead cells (and isolated nuclei) are permeable to Trypan Blue and will stain blue. Cells with an intact membrane will exclude the dye, and thus not stain. For fresh material aim for viability >90% (or as high as feasible given sample type) **[NOTE 19]**.
4. Examine cellular / nuclear morphology (and the amount of sub-cellular debris), remembering that a low-quality sample can compromise CUT&RUN.

#### 4.1.2 Cell processing

##### 4.1.2.a Fresh native cells

1. For cell cultures, aim for 70% confluency at harvest *vs*. overgrown cells.
2. Count starting material and confirm cellular integrity / morphology (**4.1.1**). Collect and process ∼20% excess cells to account for sample loss through handling and QC checks.

a. Suspension cells can be transferred directly to 15-50 mL collection tubes.
b. Adherent cells are recovered from growth vessels by gentle scraping or mild trypsin treatment.

i. Scraping: Remove growth media and replace with fresh serum-free media sufficient to cover the growing surface. Using a cell scraper, lightly rub across the culture area and pipette collected cells to a new tube. Add fresh media to rinse the plate/flask and collect any remaining cells. Visually inspect the plate/flask to ensure efficient cell harvest.
ii. Trypsin treatment: Remove growth media and gently wash cells once with PBS. Add trypsin (0.05% in PBS) sufficient to cover the growing surface. Incubate at 37°C for 5-10 mins, visually inspecting at regular intervals to ensure efficient cell dissociation. Inactivate trypsin with a 10x volume of complete media and pipette the cell suspension to a collection tube **[NOTES 20-21]**.
3. Count cells (as per **4.1.1**). Transfer number needed for CUT&RUN (500 k <u>per</u> <u>reaction</u> plus 20% excess) to a collection tube and directly continue (**4.3.2**).
4. If cells are to be used for CUT&RUN at a later time, transfer to cryopreservation media and use a slow freeze method (**4.1.3**).

##### 4.1.2.b Cell fixation (optional)

Light cross-linking (0.1% formaldehyde for 1 min) can preserve labile targets (*e.g.*, lysine acylation, phosphorylation or ubiquitination) and should be considered in such cases **[NOTES 16** and **22]**. Depending on their fragility, cross-linking can result in 25-50% cell loss so process more material accordingly. When testing new targets or antibodies with cross-linked material, always test native cells in parallel. Note that cross-linked cells can require an extra step to reverse cross-links prior to DNA purification in **Day 2** of CUT&RUN (**4.3.5**).

1. Confirm sample integrity (**4.1.1**). For cell cultures aim for 70% confluency *vs*. overgrown. This protocol can process up to 50 M cells. For > 50 M, divide material and process in parallel.
2. Suspension cells can be transferred to appropriately sized collection tubes and collected by centrifugation (600 x g for 3 mins at RT°C). Resuspend in fresh serum-free medium.
3. Adherent cells can be cross-linked directly in the culture plate/flask. Remove growth media and replace with fresh serum-free medium.
4. Add 37% formaldehyde (2.7 uL per mL of serum-free media) directly to culture to desired final concentration (0.1%) and quickly swirl to mix.
5. Incubate as desired (generally RT°C 1 min) and quench cross-linking by adding 2.5 M glycine to a final concentration of 125 mM. Quickly swirl to mix.
6. Adherent cells can now be transferred to collection tubes (as in **4.1.2a**).
7. Collect cells by centrifugation (600 x g for 3 mins at RT°C) and resuspend in fresh serum-free media. Proceed directly to CUT&RUN (**4.3**) unless:

a. If cell cryopreservation is intended, proceed to (**4.1.3**)
b. If nuclei extraction is intended, proceed to (**4.1.4**)

#### 4.1.3 Cell cryopreservation (optional)

This section describes how to process cryopreserved cells while minimizing lysis (thus avoiding a source of CUT&RUN background). Freezing allows the user to bank samples for later study and can hugely increase convenience.

1. Prepare Cryopreservation media (90% complete growth media + 10% DMSO).
2. Harvest cells. Count and confirm high viability / expected morphology (**4.1.1**).
3. Resuspend in Cryopreservation media (∼5 M cells/mL).
4. Aliquot cells to 1.5 mL Eppendorf tubes.
5. Slow freeze (-1°C/min) aliquots in an isopropanol-filled chiller at -80°C. Keep at -80°C for long term storage (duration will need to be empirically determined for each study system, though we have seen acceptable stability for a diversity of sample / target types over 12 months) [60].

**-- Thawing cells --**

6. Immediately before use, place tubes at 37°C. Work quickly to avoid cell lysis.
7. When cells are almost thawed (*i.e.*, small amount of ice still visible in tube), remove from the heat block. Gently pipette to complete thawing / ensure a mono-suspension and remove a small aliquot to count cells and examine their morphology **[NOTES 18-19]**.
8. Collect cells by centrifugation (600 x g for 3 mins at RT°C). Carefully remove and discard supernatant (while avoiding the cell pellet).
9. Resuspend cell pellet in **Wash Buffer** (105 µL per reaction) and proceed to CUT&RUN (**4.3.3**).

**Figure 3.**
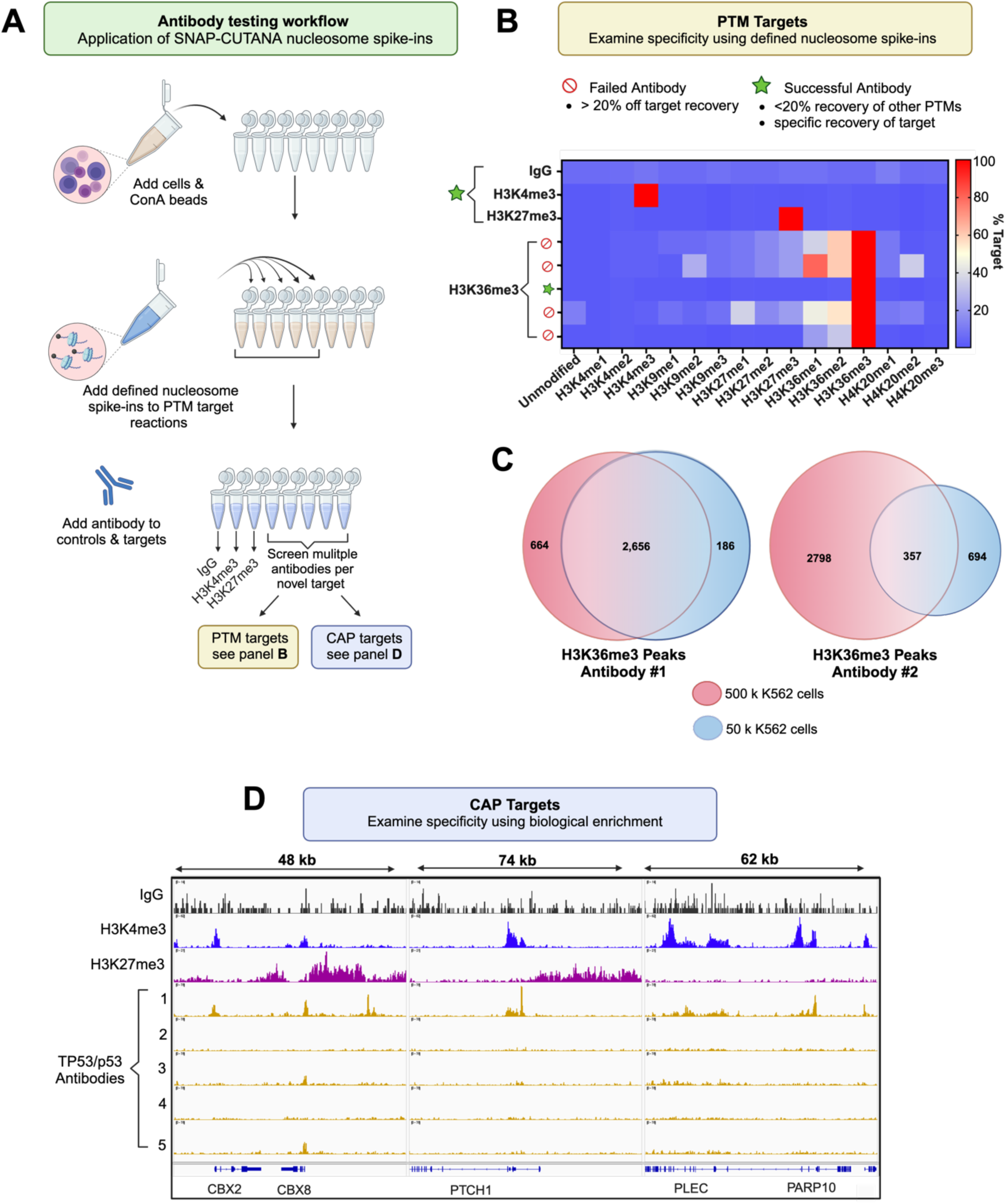
Antibody selection guidelines for robust CUT&RUN profiling^3^. **(A)** Schematic for antibody testing. Parallel reactions with cells / nuclei include controls (e.g., IgG [Negative], anti-H3K4me3 [Positive] and anti-H3K27me3 [Positive]; each with SNAP-CUTANA K-MetStat spike-in nucleosome panel) and the antibodies under test (with appropriate SNAP-CUTANA spike-in if available). **(B)** For PTMs and epitope-tagged targets, antibody specificity is determined by relative recovery of on- vs. off-target DNA-barcoded spike-in nucleosomes (see **3.3**). Representative data show pass/fail metrics for five anti-H3K36me3 tested with K-MetStat Panel (signal for each antibody by row; for each spike-in nucleosome by column). Heatmap data (key on right) reflects percentage cross-reactivity (>20% = fail) after normalization to on-target (set to 100). **(C)** Antibody efficiency in CUT&RUN is determined by peak recovery after cell dilution. When challenged by reduced cell numbers an efficient antibody (#1) retains peaks lost by a mediocre reagent (#2) (500 –> 50 k; both anti-H3K36me3 in K562 cells). **(D)** For targets without an available spike-in, multiple antibodies are compared to identify that with best balance of high yields, robust S:N and rational enrichment. In the example: of five anti-TP53/p53 tested, #1 best generates strong peak structures associated with active transcription (i.e., high correlation with H3K4me3 / low with H3K27me3). Each CUT&RUN reaction was performed with 500 k native K562 cells; RPKM normalized reads were group scaled and aligned to genome build hg38.

**Figure 4.**
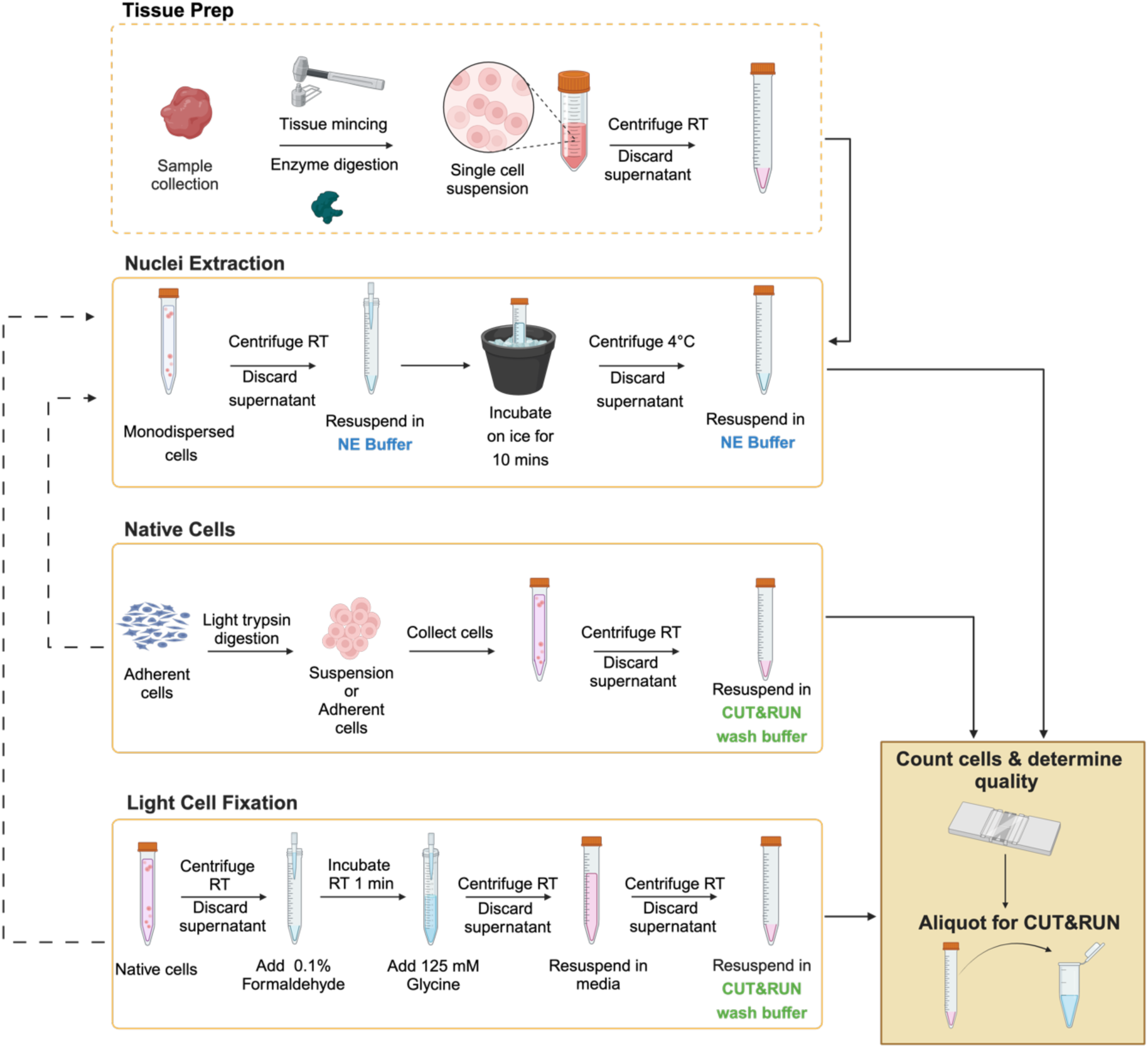
Sample preparation for CUT&RUN^4^. Schematic includes the preparation of tissues, native cells, lightly-fixed cells or nuclei for CUT&RUN **[NOTES 12-16]**. For tissues, nuclei extraction is strongly recommended. Samples are resuspended in CUT&RUN Wash Buffer (cells) or Nuclei Extraction Buffer (nuclei), counted, and checked for integrity, morphology and debris.

#### 4.1.4 Nuclei extraction (Required for tissues and many immune cells / Optional for cell lines)

Nuclei Extraction (NE) is well-suited for many sample types, but particularly tissues **[NOTES 13-15, 23]** [**Figure 4**]. If cross-linking is intended, this should be performed before nuclei are isolated. Delicate samples should be processed with an additional 25-50% cells to account for material loss. **NE Buffer** is fully compatible with CUT&RUN, and nuclei in **NE Buffer** can be directly added to activated ConA beads.

1. Prepare **NE Buffer** (2 mL per 5 M cells) fresh on the day of sample collection.
2. Count fresh or frozen cells and confirm integrity by trypan blue (or PI) staining (**4.1.1**).
3. Collect cells by centrifugation (600 x g for 3 mins at RT°C). Carefully remove and discard supernatant (while avoiding the cell pellet). Harvest 500 k cells per reaction plus ∼20% excess to account for sample loss.
4. Resuspend cell pellet in cold **NE Buffer** (100 µL per reaction).
5. Incubate on ice for 10 mins.
6. Spin cells by centrifugation (600 x g for 3 mins at RT°C). Pellet appearance should change from sticky pale-yellow cells to white fluffy nuclei. Carefully remove and discard supernatant (while avoiding the pellet).
7. Gently resuspend nuclei in cold **NE Buffer** (105 µL per reaction).
8. Remove 10 µL of sample and complete a post NE quality check by trypan blue (or PI) staining on a manual or automated counter (**4.1.1**) [**Figure 5**]. 95-99% trypan blue positive nuclei with minimal cellular debris indicates successful extraction.
9. If proceeding straight to CUT&RUN move to (**4.3.1** and **4.3.3**) of the workflow. Alternatively, nuclei can be cryopreserved for later use (**4.1.5**).

#### 4.1.5 Cryopreservation of nuclei for CUT&RUN

1. Dispense aliquots (∼5 M nuclei / mL **NE Buffer**) sufficient for intended reactions (*e.g.*, 1 mL = eight reactions plus 20% excess to account for material loss).
2. Slow freeze (-1°C/min) aliquots in an isopropanol-filled chiller at -80°C. Store at -80°C for long term storage (see **4.1.3**).
3. When ready to use an aliquot quick thaw at 37°C to minimize nuclei lysis and chromatin fragmentation.
4. Thawed nuclei in **NE Buffer** can be directly added to activated ConA beads (**4.3.3**).

### 4.2 Digitonin concentration optimization *(optional)* (day 1)

Effective cell permeabilization with digitonin is essential for a successful CUT&RUN, and it is recommended to identify the optimal detergent concentration for a new sample type. Poor permeabilization prevents antibodies and pAG-MNase from entering cells, while excess can result in cell lysis. The default digitonin concentrations in **Cell Permeabilization** and **Antibody Buffers** (0.01%) are optimal for common cell lines (*e.g.,* K562, MCF7, H1299) **[NOTE 23]**.

For new cell types, it is recommended to optimize digitonin concentrations as below. We recommend using the minimal amount that permeabilizes >95% of cells.

1. Prepare five **Digitonin Buffers**, titrating detergent concentration (0.05%, 0.01%, 0.001%, 0.0001%, 0.00001%) in **Wash Buffer**. Prepare 0.05% DMSO in **Wash Buffer** as a negative control.
2. Harvest enough cells for six CUT&RUN reactions (∼ 3.1M cells; 500 k cells per reaction plus excess).
3. Collect cells by centrifugation (600 x g for 3 mins at RT°C). Carefully remove and discard supernatant. Resuspend pellet in 620 µL RT 1X PBS.
4. Aliquot 100 µL cells to six 1.5 mL tubes. Assign one of the buffers from Step 1 to each tube.
5. Collect cells by centrifugation (600 x g for 3 mins at RT°C). Carefully remove and discard supernatant. Resuspend each cell pellet in 100 µL of the assigned buffer.
6. Incubate at RT°C for 10 mins.
7. Remove an aliquot of each sample and determine % permeabilization (aiming for >95%) with Trypan blue (**4.1.1**) or an alternate membrane impermeant dye.

### 4.3 CUT&RUN

#### ConA bead binding and antibody incubation (Day 1)

In this section of the workflow (**4.3.1** **-** **4.3.3**) ConA beads **[NOTE 24]** are activated **[NOTE 25]** and bound to prepared cells in CUT&RUN **Wash Buffer**. SNAP-CUTANA nucleosome spike-in controls are added to appropriate reactions before target-specific primary antibody is added and incubated overnight.

#### 4.3.1 ConA bead activation

1. Transfer 11 µL per planned reaction of ConA beads to a 1.5 mL tube **[NOTE 24]**.
2. Place tube on a compatible magnetic rack until slurry clears (2 - 5 min). Pipette to remove and discard supernatant. Do not disrupt pelleted beads while on magnet.
3. Remove tube from magnet and add 100 µL per reaction **Bead Activation Buffer [NOTES 25** and **27]**. Resuspend beads by pipette mixing or gentle vortexing (medium setting 5-7: used through this protocol). Do not scrape beads, as this can damage the surface coating. Flash spin to collect.
4. Return to magnet until slurry clears (2 - 5 mins). Pipette to remove and discard supernatant.
5. Repeat Steps 3 and 4 for a total of two washes. Beads should tightly collect on magnet **[NOTE 26]**.
6. Remove tube from magnet and resuspend beads in 11 µL per reaction **Bead Activation Buffer**.
7. Aliquot 10 µL activated beads into 8-strip PCR tubes. Keep tubes on ice. **[NOTE 28]**

#### 4.3.2 Initial Cell Harvest

For efficient capture by ConA beads, cells / nuclei with intact membranes should maintain morphology and be monodispersed [**Figure 1A** (step 1) **& Figure 5**] **[NOTE 29]**.

1. Count starting cells (as in **4.1.1**) and harvest 500 k per reaction (plus 20% excess). Collect cells by centrifugation (600 x g for 3 mins at RT°C). Pipette to remove supernatant, leaving enough liquid to keep the cell pellet submerged (∼ 50 µL). The pellet should be visible, and the supernatant should be clear.
2. Resuspend cells in 100 µL per reaction of RT **Wash Buffer** and gently pipette mix **[NOTE 29].**
3. Repeat the above step for a total of two washes **[NOTES 30-31]**.
4. After the final wash, resuspend cells in 105 µL per reaction **Wash Buffer**.
5. Count cells and examine morphology by Trypan Blue staining (as in **4.1.1**).
6. Capture representative sample images and proceed (to **4.3.3**).

#### 4.3.3 Sample binding to solid support

1. For each reaction add 100 µL cells or nuclei to 10 µL activated ConA beads in 8-strip tubes. When first using this protocol or working with a new sample type, process an extra reaction in parallel to directly evaluate cell integrity and effective bead-binding (*i.e.*, QC reaction) **[NOTE 32]**. Gently vortex and flash spin in a tabletop mini centrifuge to collect liquid (where ConA beads should remain dispersed).
2. Incubate sample-bead slurry at RT°C for 10 mins (where ConA beads should bind cells / nuclei).
3. Place tubes on an 8-strip tube compatible magnetic rack until slurry clears (2 - 5 mins). Remove supernatant and save 10 µL (*aka*. *unbound fraction*) into a new 1.5 mL tube.
4. Remove from the magnet. Immediately add 50 µL cold **Antibody Buffer** to each reaction (creating the *bead fraction*). Gently pipette to resuspend. If working with > 8 reactions, work with one 8-strip at a time to avoid ConA bead dry-out.
5. Transfer 10 µL of the *cell*:*bead slurry* from the QC reaction to a new 1.5 mL tube.
6. Visually inspect the *unbound fraction* and *bead fraction* on a brightfield or phase contrast microscope following the Trypan Blue staining protocol (**4.1.1**). This QC examines if sample has successfully bound to ConA beads [**Figure 1A** (step 2)]. Permeabilized cells / nuclei should appear blue (>95%) and be surrounded by beads **[NOTE 33]** [**Figure 1A** (step 2)].
7. Add **SNAP-CUTANA Spike-in Panels** to designated reactions (as applicable). Do not vortex the spike-in panel stock tube. Instead, flash spin and pipette mix prior to dispensing. If using 500 k cells per reaction, add 2 µL SNAP-CUTANA Spike-in Panel; if using <500 k cells, dilute panel appropriately **[NOTES 34-35]**.
8. Gently vortex 8-strip tubes and flash spin to collect liquid.
9. Add 0.5 µg target-specific antibody to each reaction. Pipette up and down 3-5 times to clear tip **[NOTE 36]**.
10. Gently vortex reactions to mix and flash spin to collect liquid. Incubate on a nutator overnight at 4°C with 8-strip caps elevated at 45° angle. Do not put tubes on a rotator: when rotating end-over-end ConA beads stick to the lids, leading to bead drying and sample loss.

**--- OVERNIGHT 4 °C ---**

**Figure 5.**
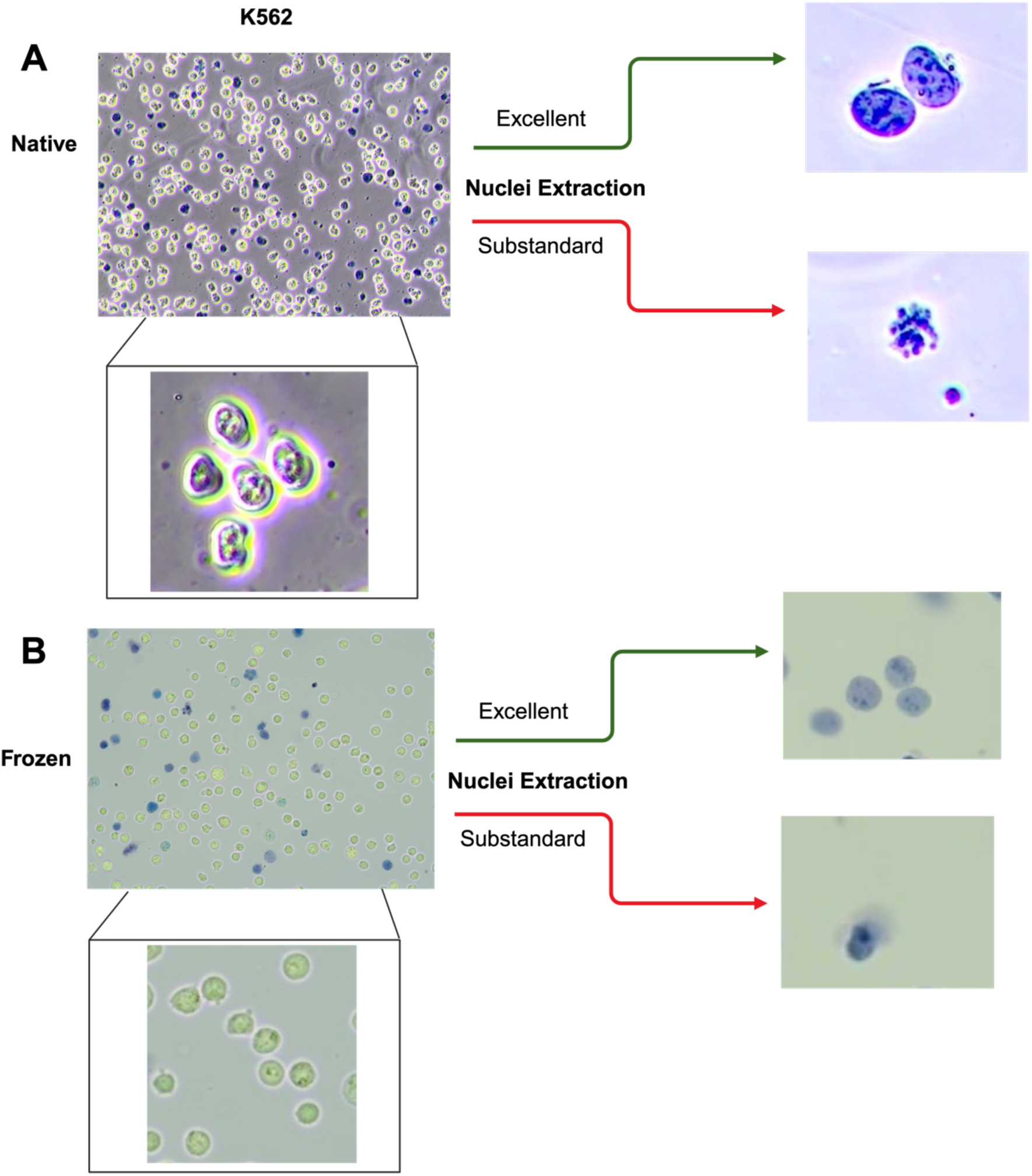
Nuclei extraction quality control^5^. Microscope images show representative morphology after nuclei extraction from **(A)** native or **(B)** Cryopreserved K562 cells. Trypan blue staining shows starting cells have high potential viability (dye exclusion) and expected morphology. Successfully extracted nuclei are Trypan blue positive with intact nuclear membranes; critical for ConA bead binding. A disrupted nuclei membrane indicates poor quality material: if the predominant species, such a sample will yield poor quality CUT&RUN data.

#### Targeted chromatin release and DNA purification (Day 2)

On protocol **Day 2** reactions are washed to remove excess primary antibody, and pAG-MNase is used to cleave and release antibody-bound chromatin fragments into solution **[NOTES 37]**. Bulk chromatin and cellular components stay bound to ConA beads and are easily removed by magnetic separation. CUT&RUN enriched DNA is then purified and readied for library preparation. Analyzing this CUT&RUN-enriched DNA on a TapeStation or Bioanalyzer will generally be uninformative, as raw yields are typically below their limit of detection. Instead, CUT&RUN DNA yields can be quantified by fluorometric assays (*e.g.*, Qubit) following manufacturers protocol.

#### 4.3.4 Chromatin Digestion

ConA beads become sticky after overnight incubation at 4°C; pipette carefully to resuspend, and regularly vortex to keep beads in solution **[NOTES 38-39]**. Ensure all liquid is ejected from pipette tips at each step to avoid bead loss. If concerned by sample loss, vortex mix reactions instead.

1. Remove reactions from 4°C and flash spin to collect liquid. Bead settling after overnight incubation is common and does not indicate experimental failure. Place on an 8-strip tube magnet until slurry clears and the beads form a tight pellet (2 - 5 min). Remove **Cell Permeabilization Buffer** from 4°C and place on ice.
2. Use a multichannel pipette to carefully remove and discard supernatant. If processing ≥8 reactions, complete buffer exchange one 8-strip at a time to avoid ConA bead dry-out.
3. Keeping tubes on the magnetic rack, immediately add 200 µL of cold **Cell Permeabilization Buffer**. Leave for ∼10-30 sec, and then pipette to remove and discard buffer.
4. Repeat for a total of two washes with **Cell Permeabilization Buffer**.
5. Prepare a master mix of pAG-MNase (2.5 µL per intended reaction) in cold **Cell Permeabilization Buffer** (50 µL per intended reaction), with extra volume to account for pipetting loss **[NOTE 38]**.
6. Remove tubes from magnet and resuspend beads in 52.5 µL cold pAG-MNase master mix per reaction. Gently vortex and/or pipette mix and flash spin to collect liquid **[NOTE 39]**.
7. Incubate reactions RT°C for 10 mins on a nutator with caps elevated at 45° angle.
8. Gently vortex tubes to mix and flash spin. Place on magnetic rack until slurry clears and the beads form a tight pellet (2 - 5 min). Pipette to remove supernatant.
9. Keeping tubes on magnet, add 200 µL cold **Cell Permeabilization Buffer** per reaction. Leave for ∼10-30 sec, and then pipette to remove and discard buffer.
10. Repeat for a total of two washes. Then place tubes on ice (using a pre-chilled aluminum block can greatly improve handling).
11. Prepare a master mix of CaCl_2_ (1 µL of 100 mM stock per intended reaction) in cold **Cell Permeabilization Buffer** (50 µL per intended reaction, with extra volume to account for pipetting loss) **[NOTE 38]**.
12. Resuspend each reaction in 51 µL cold **CaCl_2_ master mix**. Keep tubes on ice or a pre-chilled aluminum block throughout.
13. Gently vortex and/or pipette mix to ensure beads are resuspended and flash spin to collect liquid.
14. Incubate reactions 4°C for 2 hr on a nutator with caps elevated at 45° angle.

#### 4.3.5 Chromatin Release

During the CaCl_2_ incubation, prepare **Stop Buffer** and DNA purification reagents. *E. coli* spike-in DNA can be added to **Stop Buffer** as a master mix and used to monitor CUT&RUN library prep and aid sequencing normalization **[NOTE 40]**. Add 0.5 ng E. *coli* DNA per 500 k cells (scaling linearly for fewer cells).

1. Prepare **Stop Buffer master mix** by combining 1 µL *E.coli* DNA (0.5 ng/µL) and 33 µL **Stop Buffer** per reaction. *E.coli* can be added to any target reaction. Gently vortex to mix, flash spin, and place on ice **[NOTE 41]**.
2. Set a thermocycler to 37°C.
3. After the 2 hr incubation (**4.3.4** step 14), vortex and flash spin the 8-strip tubes. Add 34 µL **Stop Buffer master mix** to each reaction. Gently vortex to mix, flash spin, and place in 37°C thermocycler for 10 min **[NOTE 42]**.
4. Vortex and flash spin the 8-strip tubes. Place on 8-strip tube compatible magnet until slurry clears (30 s - 2 min).
5. Transfer supernatant to new 8-strip PCR tubes. **<u>Do not discard supernatant: this</u> <u>is the target-enriched chromatin</u>**!
6. For native materials, proceed to DNA purification (**4.3.6**). To reverse cross-links, continue below.

#### Cross-linking reversal (*optional*)

CUT&RUN cross-linking can cause cell lysis and/or reactions to become clumpy or sticky. If this occurs do not complete cross-linking reversal instead move directly to DNA purification (**4.3.6**).

7. Add 0.8 µL 10% SDS to each CUT&RUN-enriched chromatin sample. Pipette up and down 3-5 times to fully clear tips.
8. Add 1 µL of 20 µg/µL Proteinase K to each reaction. Pipette up and down 3-5 times to fully clear tips. Gently vortex to mix and flash spin.
9. Place tubes in a thermocycler at 55°C. Incubate 1 hour to overnight.

**--- *cross-link reversal* 1 HOUR or OVERNIGHT 55°C ---**

10. Flash spin tubes and proceed to DNA purification (**4.3.6**).

#### 4.3.6 DNA Purification

Many DNA purification approaches, including spin columns and magnetic beads, are compatible with the CUT&RUN workflow. However this protocol uses SPRIselect beads **[NOTE 43]** to allow streamlined DNA purification in 8-strip tubes [61].

1. Prepare 85% ethanol (EtOH) fresh 100% EtOH and molecular grade H_2_O. Allow 500 µL per reaction (includes extra volume to account for pipetting loss).
2. Thoroughly vortex SPRIselect bead reagent in manufacturer’s buffer to an even suspension (this will maximize DNA yields and reduce potential for sample loss).
3. Slowly add 119 µL SPRIselect bead reagent at 1.4X ratio **[NOTE 44]** to each sample. Ensure tips are free of bead droplets before addition **[NOTE 45]**.
4. Thoroughly pipette to mix and then vortex 8-strip on medium setting for 10-15 sec. The solution is viscous and thorough mixing is critical to good DNA yield recovery. Flash spin tubes to collect liquid and incubate at RT°C for 5 min.
5. Place tubes on a magnet until slurry clears (2 - 5 min). For washes in Steps 5-8, process one strip at a time to avoid beads drying out.
6. Carefully pipette to remove and discard supernatant, taking care not to disturb the beads. Keeping samples on the magnet, immediately add 180 µL 85% EtOH to each. Leave for ∼ 30 seconds, and then pipette to remove and discard.
7. Repeat previous step for a total of two washes.
8. Flash spin tubes to collect excess EtOH.
9. Place tubes back on magnet and remove residual EtOH with a 10 µL multichannel pipette while taking care not to disturb the beads.
10. Remove tubes from magnetic rack and air dry 2-3 mins at RT°C. Beads should remain dark brown and just barely damp **[NOTE 46]** [**Figure 1A** (step 4)].
11. Add 17 µL of **0.1X TE Buffer** to each tube. Pipette to mix beads and vortex to fully resuspend. Flash spin tubes to collect liquid.
12. To elute DNA, incubate samples 2-5 mins at RT°C.
13. Vortex tubes, flash spin, and return to the magnet. Incubate additional 2 min at RT°C.
14. Transfer 15 µL supernatant containing CUT&RUN-enriched DNA to new 8-strip PCR tubes.
15. Quantify eluted DNA with the Qubit fluorometer and 1X dsDNA HS Assay Kit **[NOTE 47].**
16. Safe point to pause: store DNA at -20°C or continue to library preparation (**4.4**).

**--- Safe pause point. Store at -20 °C ---**

### 4.4 Library Prep (Day 2 or 3)

Library prep is a multistep process whereby CUT&RUN DNA fragments are prepared for sequencing (most often paired-end on *Illumina* platforms). A range of commercial library prep kits are available (and should supply all needed reagents, including barcoded primers to distinguish each reaction library **[NOTE 48]**), but selecting one optimized for CUT&RUN (*e.g.*, **2.2**) will improve data quality and consistency.

With our preferred library prep kit 5 ng CUT&RUN DNA per library **[NOTE 49]** will provide the best balance of yield and high-quality sequencing data. If using a different kit follow its respective instructions but use the below CUT&RUN-specific PCR parameters optimized for low inputs and small DNA fragments (200-700 bp) **[NOTES 50 - 52]**. Irrespective of the kit selected, do not shear or fragment DNA prior to library prep.

| Temperature °C | Time (s) | Cycle# |
| --- | --- | --- |
| 98°C | 45 | 1 |
| 98°C | 15 | 14 |
| 60°C | 10 |  |
| 72°C | 60 | 1 |
| 4°C - 12°C | ∞ |  |

A major predictor of CUT&RUN success prior to sequencing is the <u>fragment distribution</u> of purified sequencing libraries, which should show a primary peak at ∼300 bp (∼170 bp mononucleosome + adaptors) on the Bioanalyzer or TapeStation [**Figure 1A** (step 5)]. Note that CUT&RUN library <u>yields</u> are not the only success metric, as these are influenced by target abundance, antibody efficiency, cell type and/or number.

Good libraries provide sufficient DNA to sequence at standard concentrations (>2 - 100 nM), but quality data can be generated from as low as 0.5 nM **[NOTE 52]**. If present, adapter dimers will be between 100-175 bp. If they comprise >5% of the total DNA, we recommend their removal prior to sequencing by gel purification kits [*e.g., Qiagen* 28704] or right-shifted size selection via SPRI beads (**4.5.1**).

**--- Safe pause point. Store at -20°C ---**

### 4.5 Sequencing and Data Analysis (Day 3+)

Select appropriate sequencing platform based on multiplexing requirements and desired sequencing depth. Paired-end sequencing with at least 50 bp reads (*aka*. PE100) is recommended, as this improves genomic mapping accuracy and distinguishes the nucleosome spike-in panels. CUT&RUN requires only 5-10 million total reads for appropriate peak-generation / genome coverage compared to the ∼30 million reads required by *ChIP-seq* **[NOTE 53]**. The next steps of this protocol describe how to analyze and visualize sequencing data generated from CUT&RUN libraries on a MacOS or Linux operation system.

#### 4.5.1 Pooling libraries and loading sequencer

1. Pool equimolar libraries from individual reaction libraries for multiplex sequencing.

a. Dilute each reaction library to the same concentration. 2-4 nM is recommended (if possible from available library) but confirm appropriate for the intended sequencing platform **[NOTE 54]**.
b. Pool equal volumes of each library in one tube. Ensure only unique indexes are pooled so samples can be effectively demultiplexed.
c. Dilute pool to appropriate concentration for intended sequencing platform.
2. Load the correct volume onto a temperature equilibrated sequencing cartridge for the platform of choice **[NOTE 55]**.

##### Adapter Dimer Removal (optional)

Adapter dimers can be removed from pooled sequencing libraries (at recommended 2-4 nM) using a 1:1 ratio of [library volume : SPRIselect beads]. Save ∼10-20 µL of <u>pre</u>-cleaned library pool to confirm adapter dimer removal.

a. Pool equal volumes of normalized libraries (*e.g.*, 2-5uL per) in a 1.5 mL tube.
b. Slowly add SPRIselect beads to a final ratio of 1:1. Vortex to mix. Incubate RT°C for 5 mins.
c. Place 1.5 mL tube onto compatible magnet until slurry clears (2 - 5 mins).
d. Remove supernatant. Keeping tubes on magnet, add 180 µL of 85% EtOH (freshly prepared) to tube. Leave sample in EtOH for ∼30 seconds.
e. Remove supernatant and repeat EtOH wash for a total of two washes.
f. Place samples in tabletop mini centrifuge and flash spin to collect excess EtOH.
g. Place samples back on magnet and gently remove residual EtOH with 10 µL volume multichannel pipette. Do not disturb the pellet.
h. Remove samples from magnetic rack and air dry 2-3 mins RT°C. Beads should remain dark brown and damp.
i. Add 12-15 µL of **0.1X TE Buffer** or **Sequencing Buffer** to each reaction. Pipette mix while washing the pelleted side to resuspend beads. Follow pipette mixing with a gentle vortex and flash spin.
j. Incubate RT for 2-5 mins to elute DNA.
k. Place 1.5 mL tube onto compatible magnet until slurry clears (30s - 2 mins).
l. Carefully transfer 10.5 µL supernatant to clean new 1.5 mL tube as the <u>post</u>-cleaned library pool.
m. Run <u>pre</u>- and <u>post</u>-cleaned library pools on TapeStation to confirm successful adapter dimer removal.

#### 4.5.2 SNAP-CUTANA Spike-In Analysis

Any reaction spiked with an *EpiCypher* SNAP-CUTANA nucleosome panel (*e.g.*, K-MetStat, HA Tag, or DYKDDDDK Tag **[NOTE 1]**) can be analyzed using pipeline below. The shell script searches for and counts DNA barcodes corresponding to specific dNucs. The relative on- *vs*. off-target recovery validates antibody performance and the experimental workflow, before diving into genomic alignments. Additionally, spike-in results guide any needed optimization / troubleshooting [*e.g.*, **Figure 3B**]. Since the same amount of nucleosome panel is added to each reaction, they can also be utilized as a normalization tool.

1. Download R1 & R2 paired-end sequencing files (fastq.gz) for each spiked-in reaction. Unzip and save fastq files in a new folder.
2. Download the Shell script (.sh) and corresponding analysis files (.xlsx) and save these to the folder created in **step 1**. Applicable files are on the relevant product page at *epicypher.com* [*e.g.*, SNAP-CUTANA -K-MetStat panel, #19-1002].
3. Open shell script file in a plain text editing program (*Microsoft* Word or PDF editing programs are currently incompatible). Scroll past barcode sequences and find the analysis script.
4. The script is a loop that counts the reads aligned to each DNA barcode in a given reaction, with each nucleosome spike-in usually represented by two barcodes (A & B). Create one loop per reaction (below is customizable):

a. Copy lines between # template loop begin ## and # template loop end ##.
b. Paste the loop under the last done. Paste one copy per reaction.
c. In the first loop replace Sample1_R1/2.fastq with correct R1 & R2 fastq file names for one reaction. Repeat each loop and save.
5. In Terminal, set the directory to your folder. To change directory location, type “cd” followed by a space and drag folder into the terminal. Press return.
6. Run the script in terminal. Type “sh” followed by a space and drag the .sh file from your files into Terminal to copy the location. Press return.
7. Open the panel analysis excel file and fill in the reaction names and set the relevant **on-target** in column B. The first reaction is set to the negative control (IgG) all additional reactions select a target from the drop-down menu.
8. Copy the R1 barcode read counts from the first loop in the terminal and paste into the yellow cells in the excel sheet matching the corresponding sample in **Column C**. Repeat for R2 barcodes and paste into yellow cells in **Column D**. Repeat for each loop/reaction.
9. The excel file automatically analyses the spike-in data.

a. Calculates the total reads for each DNA barcode in **Column E** (R1 + R2).
b. Calculates the total barcode read counts for each nucleosome in **Column F** (A+B).
c. Expresses total read counts for each spike-in nucleosome as a percentage of on-target read counts found in **Column G & J**. This provides on-target recovery and specificity for each antibody.
d. **Column J** displays an auto populate table (**Output Table**), with the percent recovery of each nucleosome normalized to the on-target mark. A color gradient is used to visualize recovery of each nucleosome normalized to on-target [**Figure 3**].
e. For each reaction, the percent of unique sequencing reads assigned to each spike-in can be calculated. Type the total number of unique reads in the yellow cell titled **Uniq align reads** found in **Column B.** The percent total barcode reads are added to the **Output Table.**

#### 4.5.3 *E.coli* Spike-In Analysis

In this CUT&RUN protocol, the same scaled amount of *E.coli* DNA is added to related reactions (**4.3.5**) and **[NOTE 40]**. Therefore, the spike-in can be used as a normalization tool to scale signals across reactions. Aim for 1-5% *E.coli* reads in the total sequencing reads: >10% often indicates a poor CUT&RUN reaction (*e.g.*, antibody inefficiency or poor sample permeabilization). Any reaction spiked with *E.coli* DNA can be analyzed using the pipeline below. Normalizing is valid for multiple CUT&RUN data sets generated from the same antibody target, or experimental vs. IgG negative control antibody. Applying a scaling factor across CUT&RUN data from different antibody targets (*e.g.*, H3K4me3 *vs*. H3K27me3) is invalid because there is no expectation that the *E.coli* bandwidth should be the same with epitopes of differing genomic abundance.

1. Data from demultiplexed CUT&RUN sequencing libraries requires separate alignment to the *E.coli* genome (K-12 MG1655), while filtering duplicate reads.
2. Calculate spike-in bandwidth by dividing total uniquely aligned reads to the *E.coli* genome by total uniquely aligned reads to the sample reference genome (*e.g.*,hg38).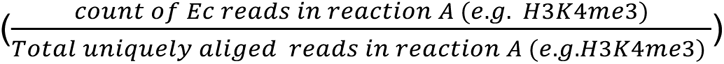
3. The normalization scaling factor is calculated such that *E.coli* spike-in signal is equal across paired reactions [62]. 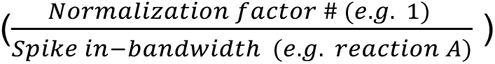
4. Use normalization scaling factor with the --scaleFactor option enabled in deeptools bamCoverage tool to generate normalized bigwig files for visualization (*e.g.*, on Integrative genome viewer (IGV)) **[NOTE 56]**.

#### 4.5.4 Genomic sequencing data analysis

CUT&RUN produces clean genomic mapping profiles using many of the same data analysis pipelines and peak calling tools developed for *ChIP-seq* (albeit with some important exceptions). Various segments of the bioinformatics pipeline transform raw sequencing reads into data for visualization and interpretation [**Figure 6**]. The initial stages of analysis involve read pre-processing, demultiplexing, sample identification and processing, and genome alignment. Within this, sequence files from individual reactions are identified by their assigned barcode, and filtered to remove duplicate reads, unaligned reads, and reads to repetitive DAC exclusion list regions. The filtered files are then processed by secondary analyses, including peak calling and motif enrichments. Below describes our recommended pipeline to analyze CUT&RUN sequencing data, using SNAP-CUTANA nucleosome and *E.coli* spike-in controls to identify assay success / quality, and troubleshooting (if needed).

**Figure 6.**
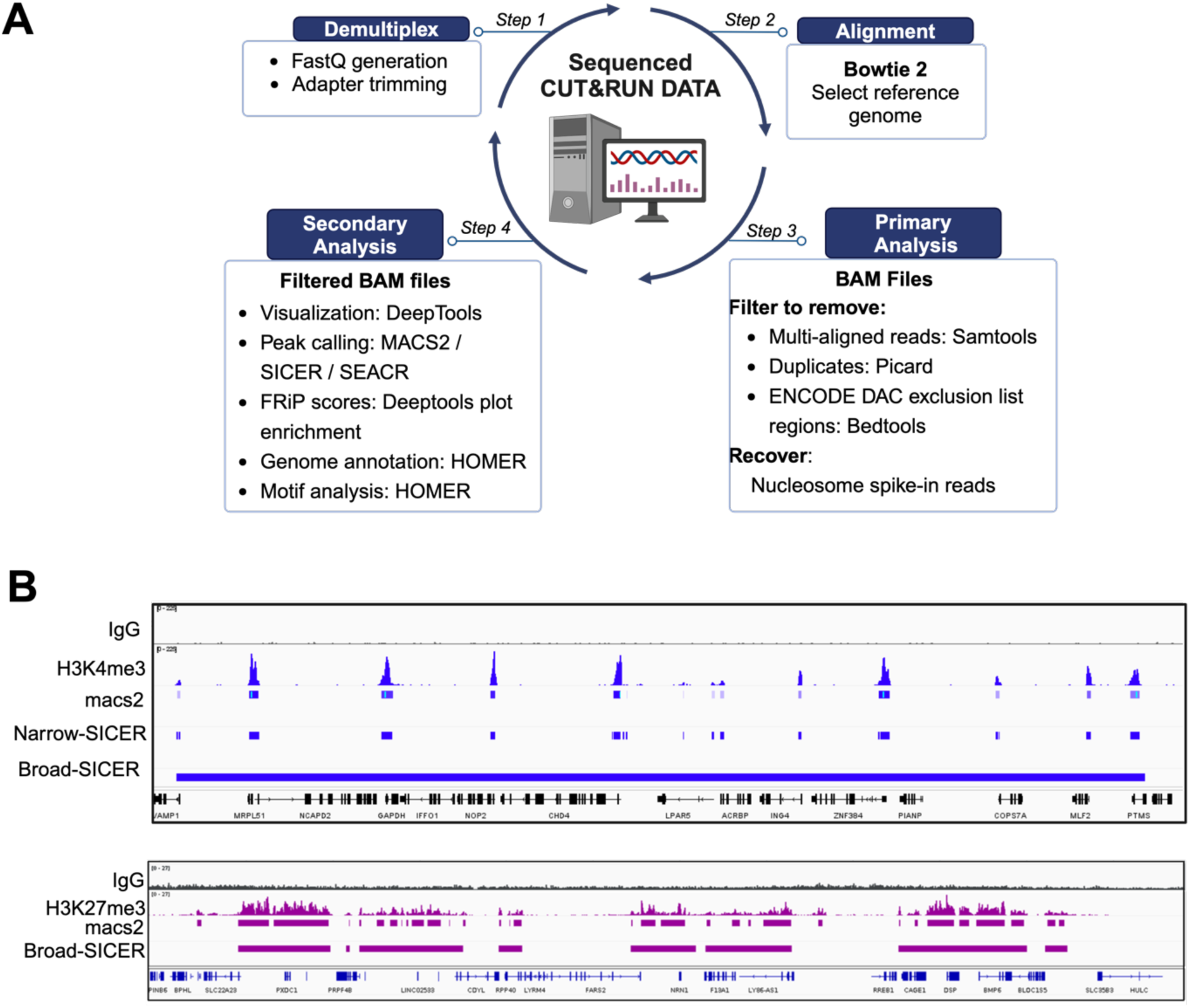
CUT&RUN data analyses pipelines and peak calling examples^6^. **(A)** Bioinformatic workflow and recommended toolkits to analyze raw fastq sequencing files from CUT&RUN experiments. **(B)** Comparison of peak-calling algorithms. MACS2 is preferred for calling targets with sharp, narrow peaks (1-2 kb; e.g., H3K4me3), while SICER is ideal for targets that enrich over broad regions (>10s of kbs; e.g., H3K27me3). All experiments used 500 k K562 cells/reaction; RPKM reads aligned to hg38 genome. H3K4me3 peaks were called by MACS2, H3K27me3 peaks were called by SICER.

Download CUT&RUN fastq files and align raw reads to reference genome of choice (*e.g.*, human hg38, mouse mm10, *Arabidopsis* TAIR10) with Bowtie2, generating a BAM file.

1. BAM files are filtered to remove duplicate reads, multi-aligned reads, and reads within DAC exclusion list regions, generating a set of **uniquely aligned reads**. Ideally >70% of total reads in a sequencing library uniquely align to the genome.
2. Filtered BAM files are converted using deepTools2 to sequence depth (and optionally spike-in) normalized enrichment tracks (Bigwig files) that enable visualization (*e.g.*, on IGV) **[NOTE 57]**.
3. Reads from SNAP-CUTANA barcoded nucleosomes or *E.coli* DNA spike-ins can be used to validate the workflow and identify / troubleshoot failed reactions (**4.5.2** and **4.5.3**).
4. Different peak ‘shapes’ (*i.e.* patterns of target localization) are called with varying efficiencies on narrow (suitable for H3K4me3) or broad (suitable for H3K27me3) settings using a variety of programs, such as MACS2, SICER, SEACR [63], or CUT&RUNTools 2.0 [64, 65] [**Figure 6B**]. It is recommended the user consider (or at least understand the pros / cons of) several peak callers and select one that best represents the target of interest [65] [**NOTE 58**].
5. For computing S:N, use software such as Deeptools plotEnrichment to calculate fractions of reads in peaks (FRiP), and compare FRiP scores from experimental *vs*. controls reactions [66] **[NOTE 59]** [**Figure 6C**].
6. Spike-in based normalization can be accomplished using reads from *E.coli* aligned reads (**4.5.3**).
7. Comparative analyses between two or more data sets (antibody or cell titrations, drug treatments, different targets, *etc*) can be done with Deeptools heat maps or Venn overlaps of called peaks [**Figure 3C**].

## NOTES

### Antibody selection

1. SNAP-CUTANA*^TM^*Spike-in Controls (*EpiCypher*) are panels of semi-synthetic, DNA-barcoded nucleosomes carrying defined PTMs or common epitope tags. Panels are bound to streptavidin magnetic beads, and added to CUT&RUN reactions just prior to antibody. During incubation, antibodies bind their target epitope in the spike-in panel, allowing targeted cleavage and release of the relevant barcoded nucleosome from bead to supernatant alongside target-enriched sample chromatin. Off-target spike-in nucleosomes remain bead-bound, and are magnetically removed alongside bead-bound cells or nuclei. After DNA purification, library prep, and NGS, the relative recovery of on- and off-target spike-in nucleosomes is determined from sequencing data by quantifying their unique DNA barcodes, offering a direct readout of antibody performance and assay success. Spike-in reads can also be used to derive a normalization standard, creating a quantifiable way to measure target-enrichment differences induced by disease or drug treatment. Available SNAP-CUTANA Spike-in Controls can be found at: https://www.epicypher.com/products/nucleosomes/snap-cutana-spike-in-controls.
2. Due to the distinct nature of immunotethering approaches *vs*. ChIP, antibodies must be empirically tested for potential adoption in CUT&RUN. In ChIP assays, bead-bound antibodies capture target fragments in the milieu of ‘sticky’ and typically denatured chromatin which drives background. This is addressed by stringent washing that can also remove on-target binding, leading to the low Signal:Noise (S:N) inherent to ChIP. Many frequently used “ChIP-grade” antibodies have been selected by the scientific community because of their high yield, which can be a product of high efficiency and/or off-target binding [67]. However, even the most specific antibodies in ChIP could require high-stringency washing to remove off-target binding, and this element is not present in the CUT&RUN workflow.
3. Testing five independent antibodies may seem excessive, but this initial investment can help to identify the best reagent, increasing confidence in all subsequent data. From our extensive testing (hundreds of targets to date), three to five antibodies are typically required to find one that works convincingly (absent a confirmed knockout). When selecting reagents, antibodies ‘validated’ for ChIP, immunofluorescence, or FACS are a good place to start but does not guarantee success. Multiple companies often distribute the same reagent, making it important to select truly independent antibodies by looking for distinct host species, immunogens, or clone identifiers. The candidate pool can include monoclonals (mAbs), polyclonals (pAbs) or recombinants, with monoclonals and recombinants often preferred to reduce (but not completely avoid) potential lot-to-lot variability. However, pAbs provide a range of paratopes and may thus be more resistant to clonal-liabilities, such as sensitivity of a PTM-specific mAb to nearby PTMs (*e.g.*, anti-H3K9me3 cannot accommodate co-incident H3S10ph) [68].
4. After comparing data from all test antibodies, decide which (if any) are suitable for further experiments based on rational pass / fail metrics: *e.g.*, % cross-reactivity to barcoded nucleosome standards **[NOTE 1]**, or independent reagents yield the same peak structure that make biological sense (such as anti-TF enriches regions containing the cognate DNA-motif). To sidestep potential future lot-variability, <u>now</u> is the time to secure sufficient material from the test lot to cover all intended studies.
5. For proteins that lack usable antibodies, epitope-tag fusions (*e.g.*, FLAG, HA, GST, or 6HIS) and related anti-tag antibodies can be explored. Epitope tags can be truly enabling but come with caveats: potentially impacting protein folding, stability or cellular localization. Further, the tag might not be accessible when the protein is part of a complex [69].
6. It is recommended to initially test and validate antibodies at 500 k cells (the standard CUT&RUN reaction) and then downscale inputs to explore antibody efficiency: when do weaker peaks drop out? If the intended study requires rare or precious cell populations, it may be useful to perform antibody validation using a relevant but more easily accessible cell line or type.
7. For CAPs, confirm the target is expressed and localized to chromatin (if feasible). Consider whether this might be impacted by conditions of cell growth and/or stimulation.
8. SNAP-CUTANA spike-in panels are added to the designated CUT&RUN reactions before target antibody. They should only be added to reactions mapping a target within the panel; *i.e.,* the K-MetStat Panel adds little discernible value to a reaction mapping lysine acetylation or a transcription factor. Reads assigned to spike-in barcodes should comprise ∼1% of total (0.1-5% is acceptable). Results are used to gauge experimental success and guide troubleshooting.
9. The negative control antibody (IgG) should show no preference for any targets in a SNAP-CUTANA panel and low genomic enrichment (potential over-digestion can be identified by a characteristic signal at the open chromatin over active promoters [defined by H3K4me3]). Positive controls (H3K4me3 and H3K27me3) should display strong enrichment for the respective on-target spike-in nucleosome with <20% off-target binding. In genomic data, positive control reactions should also show high signal to noise, as measured by the Fraction of Reads in Peaks (*i.e.*, FRiP score).
10. When control antibodies **[NOTE 9]** generate the expected spike-in percentages and genomic feature enrichments, the CUT&RUN data is reliable and the workflow validated. Alternatively, >20% off-target binding to standards and high genomic background indicates experimental issues. Sequence data from the standards can inform on failed individual reactions, of a particular target, sample, or the entire workflow.

### Sample Preparation

11. Spermidine is required in buffers because it compensates for the absence of magnesium and acts to stabilize chromatin. Buffers containing magnesium can prematurely activate pAG-MNase causing DNA degradation.
12. Although rarely observed, addition of **CUT&RUN Wash Buffer** can lyse particularly fragile cell types or samples. If a potential concern, test compatibility on a small volume of cells prior to harvesting for CUT&RUN.
13. For immune cells (*e.g.*, T, B, monocytes), there is a potential concern that binding to ConA (an immunostimulatory lectin) can cause inadvertent activation. To remove all doubt, the experimenter might want to consider using extracted nuclei instead of whole cells.
14. Tissue samples require mono-dispersion (*i.e.* to a single cell suspension) and nuclei extraction. Tissues can be processed by mechanical maceration, douncing and/or enzymatic digestion (trypsin, collagenase or dispase) [22, 70–74]. When dispersing clumps by pipetting, tips can be cut off to increase bore size and reduce the likelihood of clogging or shearing. If clumping is observed after nuclei extraction resuspend in 1X PBS.
15. Using extracted nuclei can increase S:N for chromatin-associated protein targets that are also / primarily cytoplasmic (*e.g.*, GATA3 [75]).
16. Light to moderate formaldehyde cross-linking (0.1% to 1% for one minute) can help preserve labile targets (*e.g.*, histone lysine acylations or monoubiquitylations); likely by inactivating the targeting enzymes (*e.g.*, lysine deacetylases or deubiquitinases). However, cross-linking may also lead to cell / bead clumping and loss of material. Further, heavy cross-linking (1% formaldehyde for 10 minutes; as used for ChIP) is hugely detrimental to CUT&RUN as the resulting large DNA:protein complexes are unable to diffuse across the nuclear membrane, compromising DNA yields. We have not tested other cross-linkers and so cannot comment on useful conditions. For a novel sample type / target always test native and cross-linked conditions in parallel.
17. Resuspension in **CUT&RUN Wash Buffer** can increase permeability to Trypan Blue and reduce apparent cell ‘viability’. Instead focus on total cell counts and cellular morphology / integrity. If cell counts fall and morphology deteriorates it may be better to restart with more cells, and be careful when resuspending in **CUT&RUN Wash Buffer** to avoid losing material during early wash steps.
18. Count cells immediately after adding Trypan Blue and avoid prolonged incubation. Thawed cells may have increased permeability, and thus appear to have reduced viability. As above, it is recommended to instead focus on total cell counts, cellular morphology, and the amount of contaminating cell debris. If the majority of cells appear normal and there is minimal cell lysis and debris, move forward to CUT&RUN.
19. Viability >90% is optimal but consistently lower with some primary cells, cell lines, and/or after drug treatment. Thawing cryopreserved material will result in slightly reduced viability. Reduced viability is often accompanied by reduced DNA yields and higher sequencing background but can still yield good data.
20. Trypsin will be removed in subsequent washes.
21. If used effectively, trypsinization will disaggregate clumps to monodispersed cells without damage. Optimize cell detachment by adjusting enzyme concentration and duration. Avoid extensive digestion since it can damage cells and impede binding to ConA; if extensive trypsinization is required, consider nuclei extraction.
22. Labile PTMs can be preserved by crosslinking (**4.1.2.b**) or adding appropriate inhibitors to incubation buffers: *e.g.,* protein phosphatase or KDAC inhibitor cocktails.
23. **Antibody Buffer** and **Cell Permeabilization Buffer** contain **digitonin**, a detergent to permeabilize the cell membrane. Permeabilization is NOT required when using nuclei. However, adding the detergent to **Cell Permeabilization** and **Antibody Buffers** improves bead handling (absent digitonin they can form a film on the sides of 8-strip tubes).

### ConA bead binding and antibody incubation

24. ConA beads should never be frozen or dried out, or appear black, grainy or clumped. This will / indicates damaged beads, and the CUT&RUN will fail.
25. Bead activation occurs through a series of washes that removes ConA beads from their storage buffer to facilitate its cell isolating properties based on interactions with glycoproteins on the cell / nuclei surface.
26. If working with 16 or more reactions, complete each wash with no less than 1 mL **Bead Activation Buffer.**
27. ConA bead activation requires coordination of metal ions (Calcium and magnesium) by the lectin tetramer to ensure the ConA beads can effectively bind glycoproteins on the cell / nuclei surface. ConA beads will become sticky (often manifesting as a brown residue in the pipette tip) after washing with **Bead Activation Buffer**.
28. Keep activated beads on ice until ready to combine with samples. It is advised that ConA beads be used within four hours of activation.
29. If cell clumping is observed, add 0.01% BSA and 0.01% Tween-20 to **Wash Buffer**, and monodisperse by gentle pipetting.
30. Cell loss through centrifugation, supernatant aspiration, and resuspension steps can be a major concern. This can be minimized (but not completely avoided) by longer centrifugations and careful aspiration that does not disturb the cell pellet.
31. If sample cells are ‘fragile’ and prone to lysis in **CUT&RUN Wash Buffer**, reduce to one wash step [see also **NOTE 17]**. Cell lysis increases CUT&RUN background and is thus to be avoided.
32. If cell numbers are limiting, use a small aliquot (*e.g.*,1-5 µL) of the H3K27me3 reaction for QC checks: when using a good antibody this high abundance target can spare the material.
33. At recommended volumes, ConA beads are in excess to cells; thus, observing some free-floating beads does not indicate binding failure. However, if the unbound cell / nuclei fraction is > 50% consider restarting the experiment. Ensure beads were undamaged before starting and properly activated **[NOTES 23** and **25]**.
34. SNAP-CUTANA Spike-in Panels must be added before the primary antibody.
35. If using <500 k cells/reaction, dilute spike-ins with **Antibody Buffer** for an appropriate working stock (to maintain the preferred ratio of [sample:spike-in]). This is prepared on day of experiment and used immediately.

36. Take particular care when pipetting antibodies from high viscosity glycerol storage buffers or accuracy will suffer. In the standard reaction, primary antibody (0.5 μg) is already in excess and adding more can increase background signal.

| # of cells | Panel Dilution in Antibody Buffer | Volume added to reaction |
| --- | --- | --- |
| 250 k | 1:2 | 2 μL |
| 100 k | 1:5 | 2 μL |
| 50 k | 1:10 | 2 μL |
| < 50 k | 1:10 | 2 µL |

### Chromatin Digestion and DNA Purification

37. Protein A and Protein G both bind antibody Fc domains but exhibit different binding affinities to isotypes from different species (https://assets.thermofisher.com/TFS-Assets/LSG/Application-Notes/TR0034-Ab-binding-proteins.pdf). pAG-MNase (as used in the protocol) includes both Protein A and Protein G to ensure a broader tethering compatibility for a variety of anti-chromatin antibodies.
38. Adding pAG-MNase / CaCl_2_ via a master mix minimizes experimental variability and reduces the possibility of forgetting to add a reagent to an individual reaction.
39. To allow effective reactant diffusion, ConA beads / cells should remain in suspension through pAG-MNase binding and cleavage. Beads / cells may become clumpy through incubation, but gentle pipette mixing (with tips cut off to increase bore size) can usually resolve this issue. Flash spin to settle liquid from lids before magnetic capture / opening tubes.
40. *E.coli* DNA is added to monitor library preparation / normalize sequencing data without committing excess reads, though the precise bandwidth consumed will depend on target abundance and antibody efficiency. Under optimized conditions, a negative control (IgG) or low abundance target (*e.g.*, H3K4me3) may have 10-20% *E.coli* reads, while high abundance targets (*e.g.*, H3K27me3) have 0.1-1%. *E.coli* percentages outside the usual ranges often indicate a poor CUT&RUN reaction.
41. **Stop Buffer** chelates free Ca^2+^ to quench pAG-MNase activity.
42. Samples are heated to promote chromatin release and RNase A digestion. Higher NaCl concentrations (up to 350 mM in the final eluate) may increase CUT&RUN yields.
43. SPRIselect are paramagnetic beads coated with carboxy molecules designed to reversibly bind and size-select nucleic acids. Changing the ratio of SPRIselect reagent to sample alters the size range of DNA captured. Increasing SPRIselect ratios capture smaller fragments, while decreasing ratios capture larger fragments [76, 77].
44. Multiply CUT&RUN reaction volume by ratio number for amount of SPRIselect bead reagent (*e.g.,* 84 µL reaction x 1.4 = 119 µL beads)
45. Each lot of SPRIselect beads should be tested prior to use by mixing varying bead ratios (0.8-2X) with a low range DNA ladder [*ThermoFisher* SM1213]. A passing ratio shows >90% recovery of intended target sized DNA fragments.
46. Prior to elution step (**4.3.6**) SPRIselect beads should appear slightly damp and matte brown. Glossy beads are too wet and need more time to air dry; cracked light brown beads are over-dried. Proceeding in either extreme may impact yields.
47. Low DNA yields and/or those similar to IgG (negative control) do not necessarily indicate assay failure and may simply reflect target abundance and/or antibody efficiency. For instance, with suggested antibodies (**2.2**), low abundance H3K4me3 typically yields close to IgG (∼2-5 ng from 500 k K562 cells) while high abundance H3K27me3 yields should be > than IgG under the same conditions.

### Library Preparation

48. Individual CUT&RUN libraries are prepared with 8bp barcodes at their 5 prime (i5 index) and 3 prime (i7 index) ends. For multiplexing, each individual library must be assigned a unique i5 and i7 index pair. Library prep kits will impact how many libraries can be multiplexed in a single sequencing run (*e.g.,* NEB Multiplex Oligos allows up to 24 or 96 libraries). In a given sequencing run we recommend multiplexing at least five CUT&RUN libraries to ensure index diversity. For low complexity libraries, consider including spike-in PhiX DNA based on *Illumina* recommendations.
49. For CUT&RUN reactions with yields below 5 ng DNA, use as much as possible for library preparation.
50. Making libraries from low DNA yield reactions increases susceptibility to adapter dimers. Maintain 4°C during adapter ligation setup to minimize their formation.
51. We recommend 14x PCR cycles for standard CUT&RUN reactions. If yields are low (*e.g.*, due to reduced cell inputs) this can be increased to 16x or 18x PCR cycles. However, since starting from less complexity the read duplication rate in sequencing data will likely increase.
52. If individual libraries are <1 nM, it is recommended to repeat library prep with more starting material. Alternatively, add as much as you can of lower-concentration libraries to the sequencing pool; deeper sequencing (>10 million reads) may be necessary to fully capture read diversity.

### Sequencing and Data analysis

53. Suggested numbers for sequencing depth (5-7 M total reads / 3-5 M mapped unique reads) assume human genome size. All things being equal, the suggested sequence depth for other species scales by genome size. As reference, some common genome builds: 3,100 Mb *H. sapien* (hg19, hg38); 2,500 Mb *M. musculus* (mm10); 140 Mb *D. melanogaster* (dm6); 135 Mb *A.thaliana* (TAIR10); 12 Mb *S.cerevisiae* (S288C).
54. Do not dilute libraries below 2 nM. Pooling should use 2-5 µL of normalized libraries.
55. Carefully pipet diluted libraries to avoid bubble formation that easily arise from detergent in the **Sequencing Buffer**.
56. Reactions with the highest *E.coli* spike-in bandwidth will receive the largest reduction in signal after normalization.
57. Each sample is processed and normalized by Reads Per Kilobase per Million mapped reads (RPKM). This calculation is done by Deeptools with a recommended bin size set to 20bp [59].

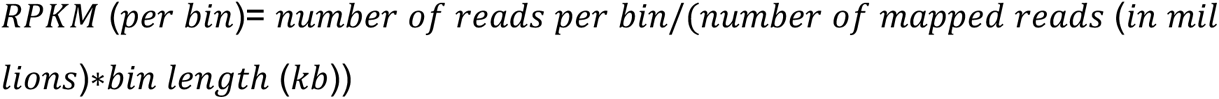
58. Various secondary analysis methods exist to identify and visualize statistically enriched narrow and broad peaks. MACS2 is a widely used peak caller that excels at detecting narrow peaks: a signal profile localized to short genomic regions usually several hundred base pairs (e.g., H3K4me3 located at promoters). SICER2 is recommended for calling broad peaks: a signal profile that covers genomic regions spanning several kilobases (e.g., H3K27me3 created by the Polycomb complex or H3K36me3 located at gene bodies). Further, the peak calling tool, and settings used, should accommodate target dynamics within a given experiment (e.g., early time-course or low-dose treatment with an EZH2 inhibitor may reduce H3K27me3 levels but paradoxically appear to increase peak numbers if the caller was optimized to join short enrichment gaps that now increase in size). If uncertain of which kind of signal profile will be generated, we recommend visualizing the data with the generated bigwig files; this will give an indication of how the profiles of diverse targets differ.
59. FRiP (Fraction of Reads in Peaks) scores reflect a peak caller understanding of S:N, where low FRiP scores identify reactions with a small number of called peaks and/or most reads outside same. This could be due to multiple technical reasons (related to antibody, sample or protocol) as described through this protocol. However, target distribution and peak caller appropriateness / settings can have a dramatic impact on FRiP scores (even in a successful experiment) so it is important to examine browser tracks and bioinformatic analyses together while interrogating the data.

## ABBREVIATIONS

CAP: Chromatin associated protein
ChIP: Chromatin ImmunoPrecipitation
CUT&RUN: Cleavage Under Targets and Release Using Nuclease
ConA: Concanavalin A
dNuc: designer nucleosome
mAb: monoclonal antibody
NE: Nuclei Extraction
NGS: Next Generation Sequencing
pAb: polyclonal antibody
pAG-MNase: protein A-protein G-Micrococcal Nuclease
RT°C: room temperature
PTM: post-translational modification
QC: quality control
S:N: Signal:Noise.

## Footnotes

1 Created in BioRender. Firestone, T. (2024) https://BioRender.com/e48e774

2 Created in BioRender. Firestone, T. (2024) https://BioRender.com/i49s432

3 Created in BioRender. Firestone, T. (2024) https://BioRender.com/n17l732

4 Created in BioRender. Firestone, T. (2024) https://BioRender.com/j87y014

5 Created in BioRender. Firestone, T. (2024) https://BioRender.com/w21q580

6 Created in BioRender. Firestone, T. (2024) https://BioRender.com/a79b961

